# Rv0233 is not essential for the survival of *Mycobacterium tuberculosis* in stress conditions

**DOI:** 10.1101/2022.03.23.485564

**Authors:** Melanie A. Ikeh, Tanya Parish

## Abstract

*Mycobacterium tuberculosis* is an important global pathogen. We were interested in understanding the role of Rv0233, a proposed subunit of the class IB ribonucleotide reductase. We constructed an in-frame, unmarked deletion strain of *M. tuberculosis* and characterized its growth and survival under replicating and non-replicating conditions. We confirmed previous studies that Rv0233 is not essential for aerobic growth or survival in the presence of nitrite. We demonstrate that the deletion of Rv0233 does not affect susceptibility to frontline tuberculosis drugs or hydrogen peroxide. The deletion strain survived equally well under nutrient starvation or in hypoxia and was not attenuated for growth in macrophages.

## Full Text

We are interested in the biology of the global human pathogen *Mycobacterium tuberculosis*, one of the largest infectious disease killers (1,2). An increased knowledge of essential processes and pathogenesis are important in identifying new drug targets and other interventions. *M. tuberculosis* has two class IB ribonucleotide reductases (RNRs); an essential RNR encoded by *nrdE* and *nrdF2* in addition to a second annotated RNR encoded by *nrdF1* and *nrdB* (Rv0233), which is dispensable for growth *in vitro* (3). Although Rv0233 was annotated as a RNR (NrdB) based on sequence similarity, this function has not been experimentally demonstrated (3,4). In addition, it was proposed that Rv0233 is not part of a functional RNR system, since it lacks the conserved C-terminal tyrosine essential for RNR function and it has homologs in other bacterial species which lack the other subunit of the class 1B RNR (4).

We constructed an in-frame, unmarked deletion of Rv0233 in *M. tuberculosis* H37Rv-LP (ATCC 25618) as described (5). Briefly, we amplified ∼1kb upstream and downstream of the gene using primer pairs MHP1-F 5’-GCT GCA GAT CTT CGG CGT ACT GGT TC-3’and MHP1-r 5’ - GAA GCT TAG GCT CGC CCA GTT GAG T-3’and MHP2-f 5’-GAA GCT TGC AGC TGG AGG ACA CCT TC-3’ and MHP2-r 5’-CGG TAC CGC TTC CAG ACG TTG CTT GTT-3’ and cloned these into the plasmid p2NIL (5). Primers were designed such that the resulting deletion would remove the majority of the 980 bp gene, leaving only a small in-frame fusion (∼150 bp). The selection marker cassette from plasmid pGOAL19 containing hygromycin resistance, LacZ and SacB (sucrose sensitivity) was added as a PacI restriction fragment (5). A deletion strain was constructed using two step homologous recombination as described (5). Briefly, the suicide delivery plasmid was electroporated into *M. tuberculosis* and single crossover (SCO) recombinants selected on hygromycin and kanamycin plates. A SCO strain was plated onto medium containing sucrose to select for double crossover (DCO) strains. PCR was used to screen colonies for the presence of the deletion or wild-type allele using primers MHP-C1 5’-CCT TAA TTA AGA CCA ACT CAG CCC AGA CC-3’ and MHP-C2 5’-CCT TAA TTA AGA TAC GCG AGT TCT GCA ACA’3’. Four potential deletion strains were analyzed by Southern blotting to confirm the chromosomal deletion (Fig 1). One strain (TAME162; Rv0233Δ) was selected for phenotypic analysis. A complemented strain was constructed by amplifying the full length Rv0233 gene using primers MHP-C1 and MHP-C2 and cloning into the integrating plasmid pAPA3 under the control of the Ag85a promoter (6). The plasmid was electroporated into the DCO strain to generate the complemented strain (TAME163; Rv0233 C’).

**Figure 1.**
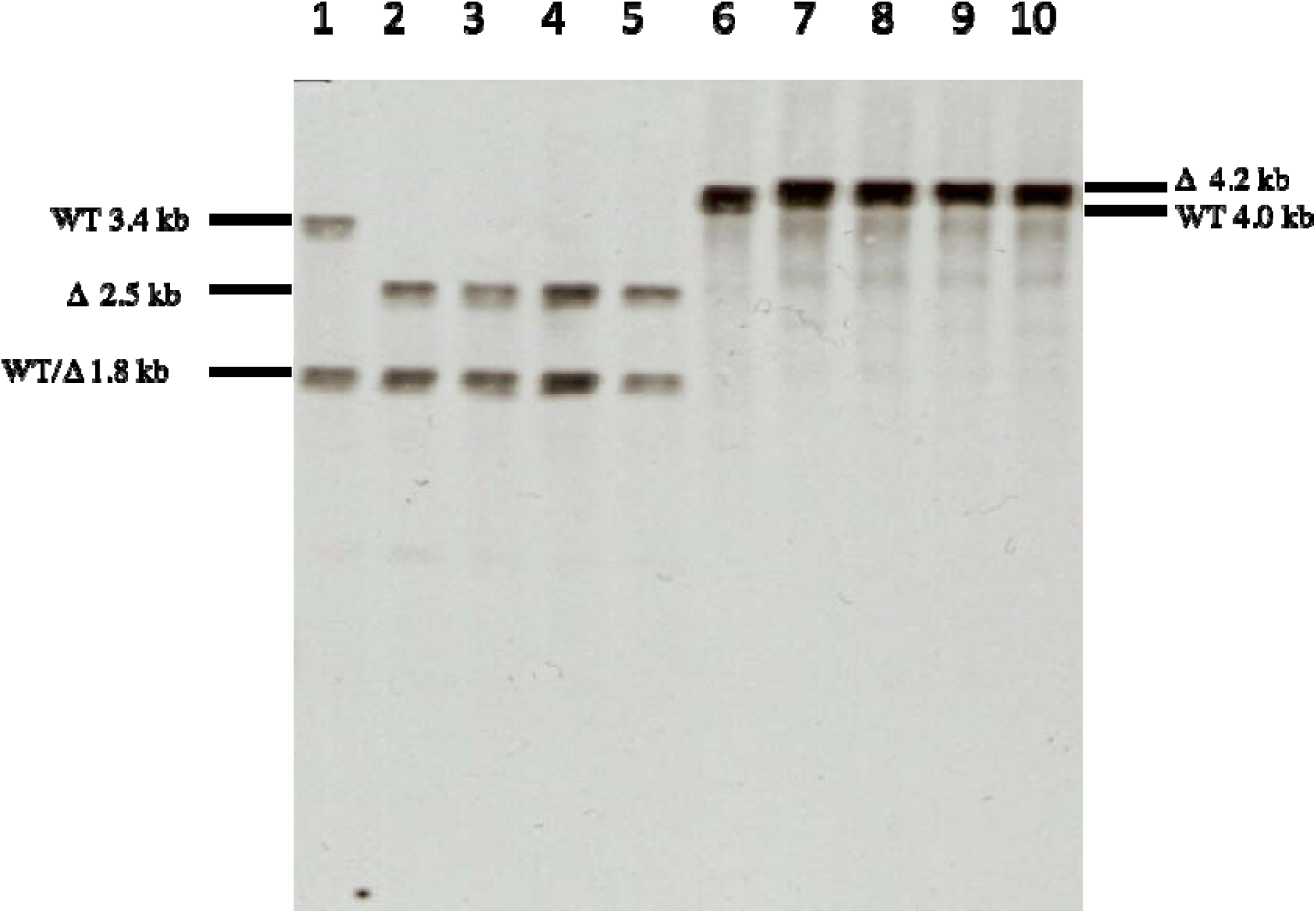
Confirmation of chromosomal deletion by Southern blotting. Genomic DNA was prepared from the wild type and four deletion strains, digested with EcoRI and KpnI (Lanes 1-5) or SacI (Lanes 6-10) and hybridized to the full length Rv0233 gene probe. Expected sizes for WT and deletion (Δ) alleles are marked. Lanes 1/6 = wild-type. Lanes 2-4 and 7-10 = Rv0233Δ strains.

Previous work had confirmed that Rv0233 was not essential and had no major effect on growth in aerobic culture (3). We confirmed this was the case for our deletion strain; where no difference in growth rate between the deletion strain, complemented strain and wild-type strain was seen (Fig 2A). Previous work had demonstrated that a Rv0233 deletion strain was equally sensitive to antibiotics that cause DNA damage or genotypic stress, but this study did not look at frontline antibiotics (3). We tested whether deletion of Rv0233 affected the minimum inhibitory concentration (MICs) of rifampicin, isoniazid, or ethambutol. MICs were determined on solid medium using the proportional method (7). There was no difference between the deletion strain and the wild-type for isoniazid or ethambutol; an increase in rifampicin sensitivity was noted, but this was small (three-fold) showing that loss of Rv0233 did not have a major impact on. sensitivity to frontline drugs (Table 1).

**Table 1.** Activity of frontline antibiotics against *M. tuberculosis* strains. Minimum inhibitory concentrations (MICs) were measured on solid medium in duplicate against wild-type and deletion (TAME162) strains. Data are mean (SD).

| | MIC ( $\mu\text{g/mL}$) | |
| --- | --- | --- |
| | Wild-type | Rv0233Δ (TAME162) |
| **Ethambutol** | 1.8 ± 1.1 | 1.8 ± 1.1 |
| **Rifampicin** | 0.063 ± 0.05 | 0.019 ± 0.01 |
| **Isonaizid** | 0.075 ± 0.04 | 0.075 ± 0.04 |

**Figure 2.**
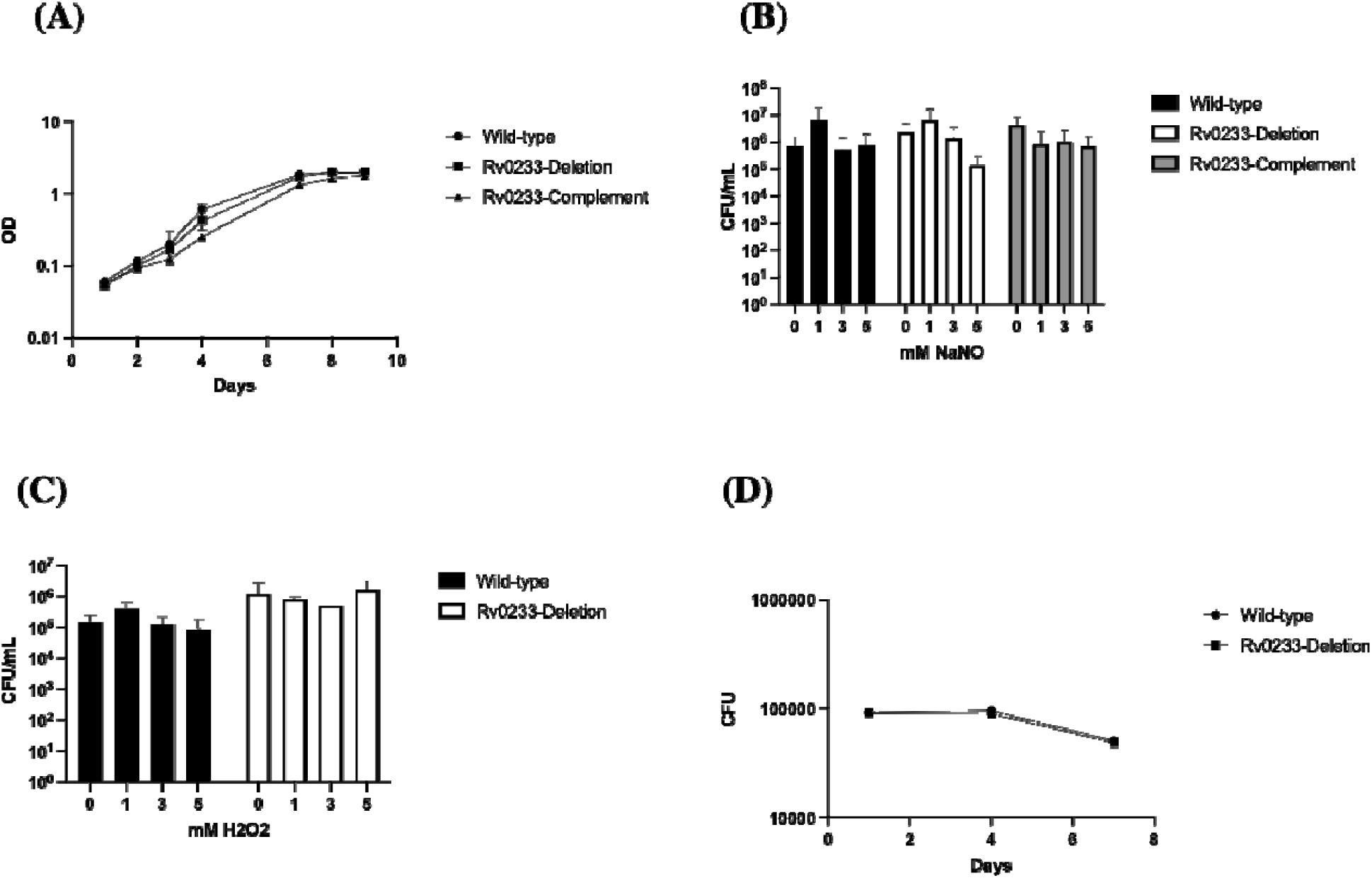
Growth and survival of *M. tuberculosis* strains. *M. tuberculosis* wild-type, deletion (TAME162), and complemented (TAME163) strains were cultured in (A) Middlebrook 7H9-Tw-OADC and growth monitored by optical density. (B) and (C) Cultures were exposed to sodium nitrite (at pH 5.6) or hydrogen peroxide for 7 days; viability was measured by CFUs. (D) Murine J774 macrophages were infected at an MOI of 1 and viable bacteria measured by CFUs. Data are the average (SD) of at least three independent cultures (n=3).

The proposed mechanism of Rv0233 predicts that it would be less sensitive to reactive nitrogen or oxygen intermediates generated by the host (4,8,9) and suggests it may play a role in surviving those conditions. We determined cellular viability after 7 days exposure to increasing concentrations of sodium nitrite (NaNO_2_ at pH 5.6) for the deletion, complement and wild-type strain and showed that there was little loss of viability in any of the strains up to 5 mM (Fig 2B). Thus, we confirmed that loss of Rv0233 had no effect on sensitivity to nitric oxide as previously reported (3). We also tested for susceptibility to reactive oxygen species; again, there was no increased susceptibility to exogenous hydrogen peroxide after 7 days treatment at increasing concentrations – the deletion strain survived as well as the wild-type strain (Fig 2C). *M. tuberculosis* is exposed to multiple stress conditions simultaneously during infection; therefore we wanted to test if Rv0233 was required to survive the hostile conditions inside the macrophage. We infected murine J774 macrophages with the deletion and wild-type strains at a multiplicity of infection (MOI) of 1:1 as described and monitored bacterial viability over 7 days (Fig 2D). We saw no difference between the deletion and wild-type strain; in both cases murine macrophages were able to control bacterial growth over 4 days with a small decrease in viable counts by day 7. Thus, the Rv0233 deletion strain was not more susceptible to macrophage-mediated killing. These data are consistent with the report that Rv0233 was not required to establish and maintain a chronic infection in the mouse model of tuberculosis (3).

Since we had confirmed that deletion of Rv0233 had no effect on replicative ability, we determined whether it was required to survive under conditions in which the bacilli are not replicating. We first tested whether the Rv0233 was able to survive under nutrient starvation. We monitored both OD and CFU over an extended period of incubation in phosphate-buffered saline (PBS) (42 days). We saw no difference in survival between the wild-type, deletion, or complemented strains, so this experiment was run a single time (Fig 3A and 3B).

**Figure 3.**
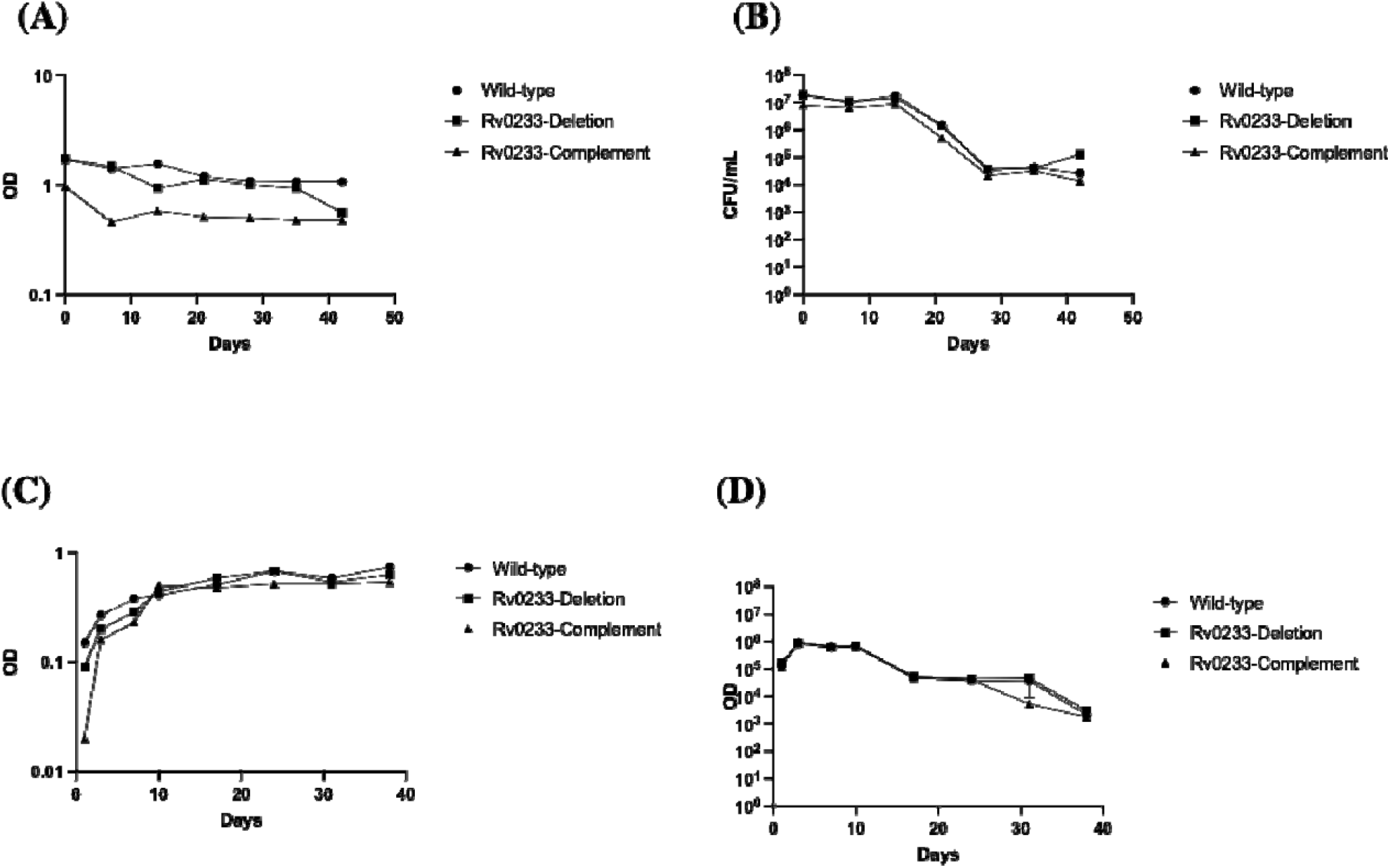
Survival of *M. tuberculosis* strains under non-replicating conditions. (A and B) *M. tuberculosis* wild-type, deletion (TAME162), and complemented (TAME163) strains were incubated in PBS. Growth and viability were monitored by OD and CFU counts. Data are from a single culture. (C and D) Strains were inoculated into DTA medium in tightly sealed glass tubes with a head space ratio of 0.5; oxygen was depleted by aerobic respiration over time. Growth and viability were monitored by OD and CFUs. Each data point is from a separate glass tube.

We next determined whether Rv0233 was required to survive under non-replicating conditions generated by hypoxia (Fig 3C and 3D). We used the Wayne model of hypoxia in which the bacteria slowly consume the available oxygen leading to a coordinated entry into a non-replicating state (10). We inoculated Dubos-Tween-Albumin (DTA) medium with each strain to a theoretical OD of 0.004 in glass tubes with a head-space ratio of 0.5. Cultures were incubated stirring to allow gradual oxygen depletion, which was confirmed by complete decolorization of methylene blue. Growth and viability were monitored by OD and CFU over 38 days. We saw no difference in survival between the wild-type, deletion, or complemented strains, so this experiment was run a single time (Fig 3C and 3D). These data show that *M. tuberculosis* does not require Rv0233 to survive under hypoxia.

In addition to confirming Rv0233 is not required for robust aerobic growth or survival during exposure to reactive nitrogen intermediates, we demonstrate that it is not required to survive oxidative stress, low oxygen, nutrient starvation and that its deletion does not make bacteria more susceptible to frontline antibiotics or macrophage-mediated killing. Since its deletion does not lead to any obvious phenotypes with relevance to survival *in vitro* or *in vivo*, or to survival of stress conditions as previously suggested (4), the role of Rv0233 in *M. tuberculosis* biology remains to be established. Taken together with its unusual structural information, we suggest that its annotation as a ribonucleotide reductase may be incorrect and that it may perform a different intracellular function, as yet not identified.

## Abbreviations

RNR: ribonucleotide reductases
SCO: single crossover
DCO: double crossover
MOI: multiplicity of infection
PBS: phosphate-buffered saline

## Author contributions

MI and TP conducted the experimental work and analyzed the data. TP wrote the first paper draft. MI and TP edited the paper.

